# Conformational Space of the Translocation Domain of Botulinum Toxin: Atomistic Modeling and Mesoscopic Description of the Coiled-Coil Helix Bundle

**DOI:** 10.1101/2024.02.02.578666

**Authors:** Alexandre Delort, Grazia Cottone, Thérèse E. Malliavin, Martin Michael Müller

## Abstract

The toxicity of botulinum multi-domain neurotoxins (BoNTs) arises from a sequence of molecular events, in which the translocation of the catalytic domain through the membrane of a neurotransmitter vesicle plays a key role. A structural study (Lam *et al.*, Nat. Comm., 2018) of the translocation domain of BoNT suggests that the interaction with the membrane is driven by the transition of an *α* helical switch towards a *β* hairpin. Atomistic simulations in conjunction with the mesoscopic *Twister* model are used to investigate the consequences of this proposition for the toxin-membrane interaction. The conformational mobilities of the domain as well as the effect of the membrane, implicitly examined by comparing water and water-ethanol solvents, lead to the conclusion that the transition of the switch modifies the internal dynamics and the effect of membrane hydrophobicity on the whole protein. The central two *α* helices, helix 1 and helix 2, forming two coiled-coil motifs, are analyzed using the *Twister* model, in which the initial deformation of the membrane by the protein is caused by the presence of local torques arising from asymmetric positions of hydrophobic residues. Different torque distributions are observed depending on the switch conformations and permit to propose an origin for the mechanism opening the membrane.

## 2 Introduction

Botulinum neurotoxins (BoNTs), secreted by *Clostrodium botulinum*, are among the most powerful toxic compounds found in nature, provoking flaccid paralysis of the host [1]. BoNTs are traditionally classified into several serotypes, termed A–G [2] and X [3]. Among them, BoNT of type A1 is the most-used BoNT in medical applications [4]. The toxins are formed by two protein chains connected by one disulfide bridge: the light chain (LC) and the heavy chain (HC). The role of HC is to prepare the delivery of the catalytic region LC which is responsible for the toxicity of BoNT. HC includes an N-terminal translocation domain (H*_N_*) (*≃*50 kDa) responsible for the LC delivery into the cytosol and a C-terminal domain (H*_C_*) (*≃*50 kDa) which recognizes specific receptors located at the terminal button of motoneurons. It is generally assumed that the two chains connected by one disulfide bridge are sufficient for producing the toxin’s physiological activity in neuromotors.

Starting from several conformations observed for BoNT/A1 [5] and BoNT/E1 [6] with various protonation schemes depending on physiological pH values of 4.7 and 7, a recent molecular modeling study [7] revealed that the global conformational variability of the toxin mainly depends on the relative displacements of the toxin domains. In addition, the analysis of the relative accessible surfaces according to pH variations, pointed to the translocation domain as one of the hotspots for the interaction between membrane and toxin in accordance to previous models [1].

The structures of the *isolated* translocation domain of BoNT/A1 [8] were determined experimentally (Fig 1A) at two pH values: 5.1 and 8.5, which are close to the physiological pH values (acidic and neutral) regulating the function of BoNT. For the acidic pH value, a dimer of two translocation domains was observed (PDB entry: 6DKK), whereas the other structure (PDB entry: 6MHJ) was monomeric and quite similar to the structure observed within complete toxin structures. In the dimeric structure, the switch domain (residues 620-666 in 6MHJ) undergoes a transition from an *α*-helix bundle to a *β* strand. The monomer *β* strands interact with each other, producing a larger *β* strand which stabilizes the inter-monomer interaction. Lam *et al.* [8] concluded from their observations that the dimerization is not relevant for BoNT/A1 function *in vivo* but that the acidification of the inner part of the neurotransmitter vesicle will induce a conformational change of the switch favoring the interaction of the translocation domain with the membrane.

**Figure 1:**
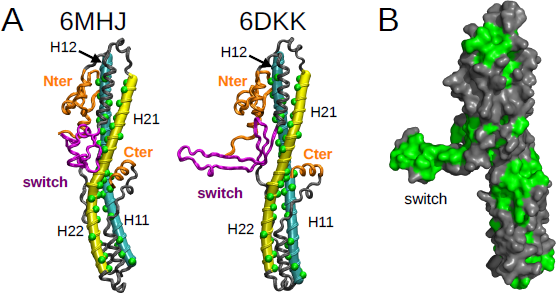
A. X-ray crystallographic structures determined for the translocation domain (PDB entries: 6MHJ determined at pH 8.5 and 6DKK determined at pH 5.1 [8]. The tubes generated by Bendix [28] on the helices 1 (cyan) and 2 (yellow) are shown for the two PDB structures. The hydrophobic residues in these helices are represented by green spheres. The limits of the helices are defined as in Table 1. The switch is colored in magenta, and the N terminal domain and the C terminal *α* helix in orange. In helices 1 and 2, the helix halves H11, H12, H21 and H22 are indicated with labels. B. Structure 6DKK represented as a surface. The hydrophobic residues GLY, ALA, VAL, LEU, ILE, PRO, PHE, MET and TRP are colored in green. This image was prepared using pymol [56].

**Table 1:** Definition of protein domains in the translocation domain H*_N_*. Differences between the two structures are highlighted in bold face. The regions H11 and H12 correspond to parts of the helix 1 in interaction with the regions H21 and H22 of the helix 2 through the coiled-coil motifs (see Section 3.3).

| Domain | Residues range 6MHJ | Residues range 6DKK |
| --- | --- | --- |
| N terminal | 547-596 | 547-596 |
| switch | 620-666 | 620-666 |
| switch tail | 663-677 | 663-677 |
| helix 1 | **692**-753 | **688**-753 |
| H11 | **692-716** | **688-714** |
| H12 | 724-753 | 724-753 |
| helix 2 | 765-**823** | 765-**827** |
| H21 | 765-**794** | 765-**793** |
| H22 | **804-823** | **800-827** |
| C terminal tail | 846-857 | 846-857 |
| C terminal $\alpha$ helix | 860-**871** | 860-**870** |

The present work intends to investigate the internal dynamics of the translocation domain in water and in a mixture of water and ethanol with the help of atomistic molecular dynamics (MD) simulations on the hundred of nanosecond time scale. Ethanol has been chosen as co-solvent for the following reasons. As the OH group of ethanol is inherently polar, while the CH3 group of its short carbon chain exhibits hydrophobic characteristics, ethanol has the ability to engage with proteins through both hydrophilic and hydrophobic interactions. In this context, ethanol has frequently been regarded as a model for investigating the equilibrium between hydrophobic interactions and hydrogen bonds. Last but most important for the present study, ethanol water solutions are usually exploited to mimic the environment of cell membranes and in particular the membrane-water interface [9, 10], providing access to a diversity of conformations for membrane proteins. A 50-50 water-ethanol ratio is a good compromise to set the ethanol content high enough to force a protein to expose its hydrophobic segments [11] while preventing the aggregation of ethanol molecules, which may impede the weakening of the hydrophobic protein core [12, 13, 14].

The starting points of the molecular dynamics trajectories will be provided by the X-ray crystallographic structures determined by Lam *et al.* [8] using monomeric conformations. Moreover, we will discuss how the protein can deform the membrane after making close contact using a biophysical model. In the past, such models have successfully explained the formation of lipid membrane rafts [15, 16], interface-mediated interactions between membrane-bound particles leading to endo-/exocytosis [17] or the formation of membrane channels [18]. To understand how an object containing hydrophobic and hydrophilic parts such as a biofilament or a protein domain can interact with a membrane, mesoscopic models are of particular interest. A combination of elasticity theory and geometry helps to describe the various system configurations. Within this framework, two mechanisms have been proposed which can induce the deformation of a membrane due to an object: the *Twister* and the *Darboux torque* mechanism [19]. Since then, the *Darboux torque* mechanism has been found to explain membrane deformation by FtzsZ [20] and by ESCRT-III [21, 22] or membrane fission by dynamin [23]. To the best of our knowledge the *Twister* mechanism was not put in evidence for explaining biological interactions with the membrane and we will explore its applicability in the framework of the BoNT-membrane interaction.

The originality of the work not only lies in the possible emergence of the *Twister* mechanism but also in the implicit modeling of the membrane in the simulation. This choice was motivated by the hypothesis that the inertia of membrane atomistic or coarse-grained systems considerably slows down the events related to the interaction between protein and membrane. The use of a mixed solvent mimicking the hydrophobic interactions produced by the membrane allows to confirm the model of anchoring of the hairpin in the membrane. The final aim of the present work is to obtain information on the protein’s conformational changes at the molecular level, in particular during the interaction with the vesicle membrane, in order to propose mechanistic hypotheses for the translocation of the full toxin through this membrane.

## 3 Results

In this section, we present the results obtained from the protein’s structure and dynamics along the MD trajectories recorded in water and in a 50-50 water-ethanol mixture. We start with an analysis of the internal mobility of the domain, followed by a presentation of the distribution of the solvent molecules. In a final step the *Twister* model is applied on the helices 1 and 2.

In the following, we selected values of 4.7 and 7 for acidic and neutral conditions, respectively, although the structures 6MHJ and 6DKK were determined at pH values of 8.5 and 5.1. These values were also those selected to mimic the acidic and neutral conditions inside the vesicles, in our previous full length BoNT MD study [7] as this allows to directly compare the behaviors of the translocation domain and of the full-length proteins. As in the present study the pH effect is modeled through different protonation patterns among protein residues, we checked, the differences in protein protonation for pH 4.7 vs 5.1 and pH 7 vs. 8.5 using H++, a method for pK_a_ calculations based on a continuum electrostatic model [24, 25] (Table 2). The following protonated residues are predicted at acidic pH: glutamic (GLU) and aspartic (ASP) acids, both protonated on the sidechain carboxyl group. The protonation of one histidine is predicted at pH 8.5. At pH of 4.7, eight residues are detected in 6MHJ, and six in 6DKK. Among them, four residues are located in the switch, one residue in the switch tail and one residue (GLU-809) in the helix 2. The three residues GLU-620, GLU-809 and ASP-848 are protonated in both structures, and for the two pH values 4.7 and 5.1. The protonation at pH=7 or 8.5 are quite equivalent for both 6DKK and 6MHJ. As for acidic pHs, there is a slight similar variation in the protonation between 6DKK and 6MHJ, as few more residues are protonated at pH 4.7 with respect to 5.1.

**Table 2:** List of protonated residues according to the studied system. These residues correspond to glutamate protonated on sidechain carboxyl (GLU/E), to aspartic acid protonated on sidechain carboxyl (ASP/D) and to one histidine protonated on H*ɛ*2. The residues protonated in both structures at a given pH value are written in bold, and the corresponding regions defined for each structure in Table 1 are given in parentheses.

| 6MHJ (pH values) |  |  |  |
| --- | --- | --- | --- |
| 4.7 | 5.1 | 7.0 | 8.5 |
| GLU-558 (N terminal),<br>**GLU-617**,<br>**GLU-620** (switch),<br>**GLU-809** (helix 2),<br>ASP-625 (switch),<br>ASP-629 (switch),<br>GLU-758,<br>**ASP-848** (C terminal tail) | **GLU-620** (switch),<br>**GLU-809** (helix2),<br>ASP-625 (switch),<br>**ASP-848** (C terminal tail) | - | HIS-561 (N terminal) |
| 6DKK (pH values) |  |  |  |
| 4.7 | 5.1 | 7.0 | 8.5 |
| **GLU-617**,<br>**GLU-620** (switch),<br>GLU-670 (switch tail),<br>**GLU-809** (helix 2),<br>ASP-616 (switch),<br>**ASP-848** (C terminal tail) | GLU-617,<br>**GLU-620** (switch),<br>**GLU-809** (helix2),<br>**ASP-848** (C terminal tail) | - | - |

### 3.1 Internal mobility of the translocation domain

The Root Mean Square Deviation (RMSD) between atomic coordinates of conformations is utilized to get global information on tertiary rearrangements in the protein with time. Atomic Root Mean Square Fluctuations (RMSF) are analyzed to investigate local motions in distinct protein regions, while the radius of a cylinder encompassing the protein structure is monitored to complement global structural results. The combined analysis of RMSD, RMSF and cylinder radius provides useful insights for tertiary modifications in different conformational and protonation states. The RMSD values were calculated from the atomic coordinates of the initial frame of the simulation, obtained from the PDB structure. The RMSF have then been calculated with respect to the average conformation along the trajectories (Table S1). The coordinate RMSD for the backbone’s heavy atoms along the MD trajectories in water (Fig 2, left panels) display different behaviours for trajectories starting from the momomer (6MHJ) compared to a monomer extracted from the dimer (6DKK). The monomeric structure 6MHJ stays stable with a RMSD plateau around 3 Å, attained after 50 ns of trajectory, whereas the monomer extracted from the dimer structure 6DKK exhibits more heterogeneous RMSD profiles along the trajectories with plateaus lying between 4 and 8 Å. For a given starting structure and a given solvent, the variation of the protonation does not induce different trends in RMSD variations. However, comparing the global RMSD values for different solvents, the values observed for 6MHJ in water-ethanol increase slightly with respect to the situation in water (Fig 2, upper four panels) whereas the values obtained for 6DKK are reduced (Fig 2, lower four panels). The water-ethanol mixture thus seems to produce opposite effects on the two conformations.

**Figure 2:**
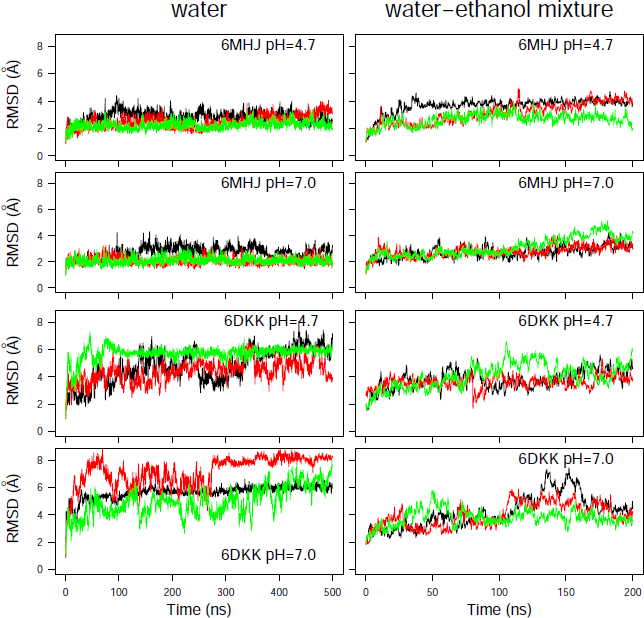
Coordinate root-mean-square deviation (RMSD, Å) along the molecular dynamics (MD) trajectories recorded in water and in 50/50 water-ethanol mixture at the two pH values. Different colors correspond to different replicas.

These differences observed for the water-ethanol mixture may be put in relation with the physiological effect of pH on the protein, as the two starting conformations were experimentally determined at different pH values. On the one hand, the structure 6DKK was obtained at acidic pH corresponding to the physiological conditions during the toxin’s interaction with the membrane of the synaptic vesicle. Since the water-ethanol mixture intends to mimic the hydrophobic effect of the membrane, it is interesting to see that the coordinate RMSD is reduced in the mixture with respect to the observations in water. On the other hand, the structure 6MHJ was observed at neutral pH, corresponding to physiological conditions of non-interaction with the membrane. Therefore, it is not surprising that the water-ethanol mixture induces a slightly larger drift of coordinates than the water solvent.

Coordinate RMSD calculated for the helices 1 and 2, as well as for the switch, are displayed as boxplots (Fig S1). The helix RMSD are mostly located in the 1-2 Å range, in agreement to the conservation of the secondary structure of the *α* helices. Nevertheless, the RMSD values for some 6MHJ replicas are larger than 2 Å in the water-ethanol mixture. In particular, the 6MHJ *α* helices exhibit RMSD values in the solvent mixture, which are slightly larger than the ones observed in water, especially for helix 2. The hydrophobic field produced by the mixed solvent thus seems to have a destabilizing effect on the structure 6MHJ. This agrees with the assignment of the structure 6MHJ to the state of the translocation domain in solution. The destabilization propagates also to the switch for one 6MHJ trajectory at pH 7 in water-ethanol solvent (see the orange box at the bottom right of Fig S1). But, almost no effect of the pH is observed in the helices of 6DKK, both in water and in ethanol-water mixture. This agrees with the observation of Ref. [26] according to which the secondary structure of the helices does not change with the pH.

In contrast to helices 1 and 2, large RMSD values are observed for the switch in most of the trajectories recorded for 6DKK. These values are larger (in the 5-8 Å range) in water than in the water-ethanol mixture (3-6 Å range). This dispersion recalls the large RMSD drifts observed for 6DKK in water (Fig 2), elucidating that the global conformational drift of the translocation domain is mostly due to the switch region.

The atomic root-mean-square fluctuation (RMSF) profiles (Fig 3) display a large peak located in the region of residues 620-660 corresponding to the switch. This peak is much smaller for 6MHJ than for 6DKK. Several smaller peaks are observed in the N terminal region, as well as in the loops connecting the helices 1 and 2 and the remaining part of the structure. For some trajectories of 6MHJ, a reduction of the pH in water induces the appearance of larger peaks in the switch as well as in the N terminal region (upper left panel of Fig 3). This gives a hint about the initial transition steps of 6MHJ towards the 6DKK conformation under acidic pH. Large RMSF peaks are also observed for the C terminal *α* helix. Looking at the conformational changes in the last frame of the trajectories, the C terminal *α* helix, undergoes unfolding in the trajectories labeled with ‘u’ (Fig S2), populating conformations similar to those observed in the equivalent region of the X-ray crystallographic structure of BoNT/E1 [6]. In other trajectories labeled ‘s’ (Fig S2), the C terminal *α* helix swings: in the full toxin, this motion would push the receptor binding domain further away from the translocation domain. This separation of the translocation and the receptor binding domains has been observed to a more limited extent during the MD simulations recorded on the full toxins [7].

**Figure 3:**
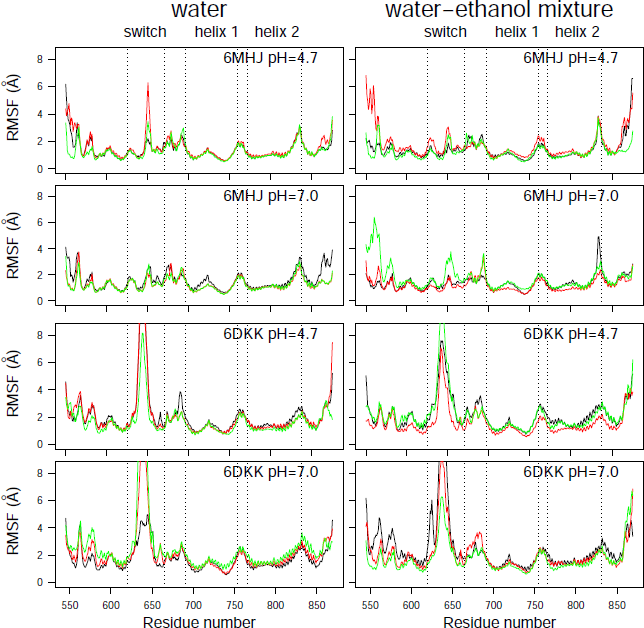
Coordinate root-mean-square fluctuations (RMSF, Å) calculated on the MD trajectories recorded on the two protein systems. Different colors correspond to different replicas.

As it is visible for the conformations displayed in Fig S2, the switch shows various orientations with respect to the remaining part of the domain. In order to get a finer picture of the switch positions, the geometry of the protein conformation was modelled as a cylinder as described in Section 5.3. The time behaviour of the cylinder radius (Fig 4) parallels the one of the global RMSD (Fig 2). The radius is quite stable around 14 Å for all trajectories recorded on 6MHJ (four top panels of Fig 4). Interestingly, the trajectories display a slight drift at acidic pH (top left panel, green curve) as well as in the water-ethanol mixture, at all pH values. For 6DKK, very heterogeneous profiles are observed for the radius with values in the range of 14.5-19 Å (four bottom panels of Fig 4). Most of the profiles display a decrease in the radius values in water along the trajectory time (bottom left panels). By contrast, the use of the water-ethanol mixture prevents the radius to decrease (bottom right panels). Indeed, the cylinder radius fluctuates around 18 Å, which is the initial radius value arising from the protruding hairpin observed in the PBD entry. This suggests that, in contrast to water, the hairpin keeps protruding from the structure in the water-ethanol mixture, as can be confirmed by visual inspection of the final system snapshots along the trajectories (Fig S2).

**Figure 4:**
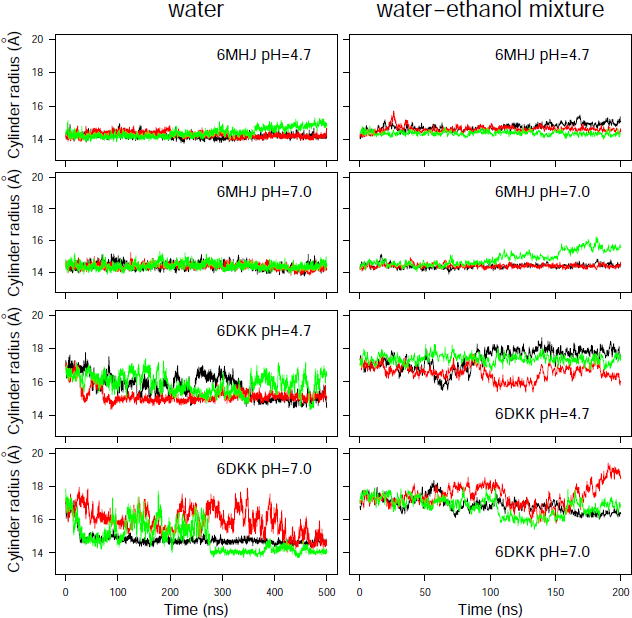
Cylinder radius *r* calculated as: 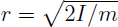 where *I* is the moment of inertia with respect to the axis of the cylinder and *m* is the total molar mass of the C*α* atoms. Different colors correspond to different replicas.

Based on all variations observed along the trajectories in water, the strong conformational drift in 6DKK (Fig 2) can be attributed to a variation of the conformation of the switch, due to a tendency to fold onto the remaining part of the translocation domain (Fig S2). Interestingly, the effect is weaker at acidic pH (see conformations in the upper left panel for 6DKK in Fig S2). This folding is due to the large hydrophobic patch present on the switch, which favors the interactions with other hydrophobic patches on the surface of the translocation domain (Fig 1B). The drift agrees with the importance of the dimer interface to stabilize the protruding conformation of the switch in the 6DKK structure and confirms that the conformation of isolated 6DKK is probably not relevant in water. At the contrary, in the water-ethanol mixture, the hydrophobic field arising from the presence of ethanol permits to keep the extended conformations and to support the model of membrane interaction proposed in Ref. [8].

### 3.2 Distribution of ethanol and water molecules around the translocation domain

To get insight in the solvent partition close to the protein, the average Spatial Density Functions (SDF) around the two protein structures in the water-ethanol mixture are calculated and the results shown in Fig 5.

**Figure 5:**
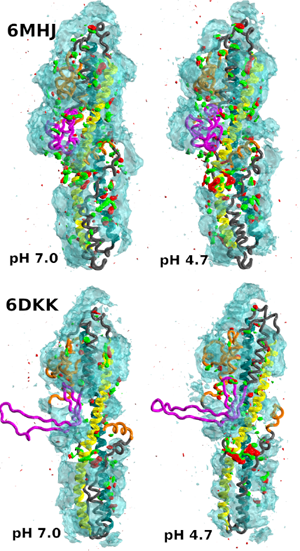
Isosurfaces of the Spatial Density Function of water and ethanol atoms around the protein. Cyan isosurface: water oxygen atoms; red and green: oxygen and methyl carbon atoms, respectively. The isosurfaces are represented at the same isodensity level (0.0115) for both water and ethanol. The protein regions are colored as in Fig 1. Data were collected from the three replica trajectories. This image was prepared using VMD [51].

Overall, the protein is preferentially hydrated. However, ethanol molecules are able to establish interactions with the protein, replacing water. When the solute largely fluctuates, the resulting SDF of the solvent molecules is averaged to zero. For this reason, the protruding switch conformation in 6DKK displays paradoxically no persistent interaction with ethanol molecules (Fig 5, bottom row). The pH acidification induces a larger number of ethanol molecules to localize in the central region. This agrees with the tendency of the translocation domain to interact with the membrane at the same pH range. Preferential ethanol solvation is, however, more pronounced in the central region of 6MHJ (Fig 5, upper row) than of 6DKK.

In Fig S3 a more detailed comparison of the solvent distributions at the two pH values is shown for 6MHJ only. Preferential ethanol solvation is observed in particular around stretches of hydrophobic residues present in the switch tail and the C terminal tail (Table 1), and highlighted in dark grey surface representation. The switch tail, located at the C terminal of the switch, corresponds to the very hydrophobic primary sequence ILLEFIPEIAIPVLG, whereas the primary sequence STDIPFQLSKYV of the C terminal tail is less hydrophobic. It is interesting to note that in the protein structure, the switch tail is closer to the protruding switch and possibly to the membrane than the C terminal tail. The persistent ethanol molecules observed close to the tail regions support a model of protein-membrane interaction in which the tails will be solubilized early during the process.

This solubilization together with the mobility and unfolding tendency of the C terminal *α* helix (Fig S2) and the mobility peaks in several RMSF profiles for the N terminal domain (Fig 3) suggests that the bundle formed by the two helices 1 and 2 will constitute the remaining piece of the folded structure involved in the interaction between the membrane and the translocation domain. This interaction will be analyzed in the following with the help of the *Twister* model.

### 3.3 *Twister* model as a possible route for opening the vesicle membrane

To apply a local force or torque distribution on a membrane and consequently induce a deformation, forces and torques have to be balanced globally at equilibrium. For a straight filament the *Twister* model suggests that a mismatch between the filament’s internal twist and the hydrophobic regions of the membrane leads to a local torque [19]. The simplest case which cancels the total torque consists of two antiparallel point torques (called the *Twister* in the limit of vanishing distance between the two torques). In the following we will study whether this model is also relevant for the case of the helices 1 and 2 in the BoNT translocation domain.

For applying the *Twister* model, we need to be sure that the *α* helix structures are stabilized at least for the first steps of the interaction with the membrane. This stability is induced by the interaction between the upper and lower halves of the helices observed in the X-ray crystallographic structures (Fig 1A). To support this point, the residues of the regions in contact, 695-713/801-819 and 730-748/766-784, extracted from the PDB entries 6MHJ and 6DKK, were processed using the software CCCP (Coiled-coil Crick Parameterization) [27], available at the Web server: www.grigoryanlab.org/cccp. This analysis (Table S2) shows that the regions in contact display coiled-coil structural motifs, stabilizing the helix conformations and consequently preserving the predefined asymmetry in the positions of hydrophobic residues. The presence of these coiled-coil motifs will indeed permit the application of local torques during the helix/membrane interaction.

An analysis of the torques *T̃_kl_*, introduced in Sec. 5.4, shows that the distribution of the hydrophobic residues along the helices 1 and 2 can induce local torques of mostly alternating sign, reminiscent of *Twister* -like interactions (see Figs. 6 and S4, S5, S6, S7, S8). The configurations of the helices are stable, which is visible in the coordinates RMSD smaller than 2 Å, displayed in Fig S1 and reflected by the tiny standard deviations of *T̃_kl_*. The only exception is observed for a residue located in the middle of helix 1, ALA-719, which is in close spatial proximity to the switch.

**Figure 6.**
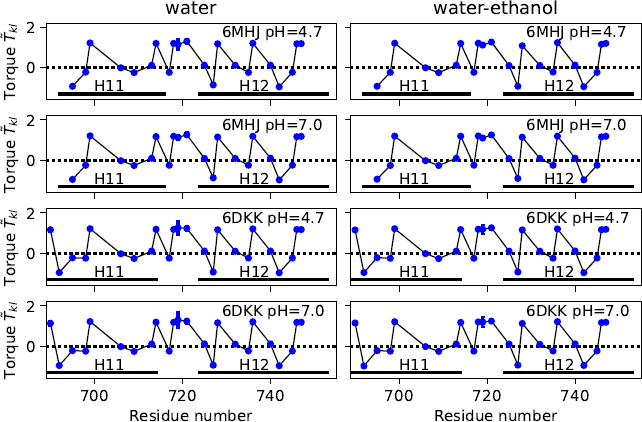
Local torque values *T̃_kl_* along the helix 1. Values averaged along the first trajectory replica and their standard deviations are plotted as a function of the residue number. Standard deviations are smaller than the size of the data points except for residue ALA-719.

In order to investigate more deeply the configuration of the helices, we have used the visualisation processing Bendix proposed for protein *α* helices [28]. This processing provides additional information for the helix coarse-graining: first, it shows that a representation as elastic tubes is applicable and second, it allows to calculate the local bending angles along these tubes. The bending angles along the helices 1 and 2 (Figs 7, S9 and S10) display similar profiles in all conditions. The profiles are almost superimposed for helix 1, whereas small variations with pH and type of solvent are observed for helix 2. A greater sensitivity to the variation of conditions had already been shown for helix 2 in 6MHJ in the subdomain RMSD (Fig S1). The large peaks observed at the middle of the profiles denote a singularity in the definition of the helix axis, implying a kink. This again suggests to consider two parts of each helix separately in agreement to the previous coiled-coil analysis. We propose thus to define four helix parts: H11, H12 for helix 1 and H21, H22 for helix 2, with the residues ranges given in Table 1 (see also Fig. 1A).

**Figure 7:**
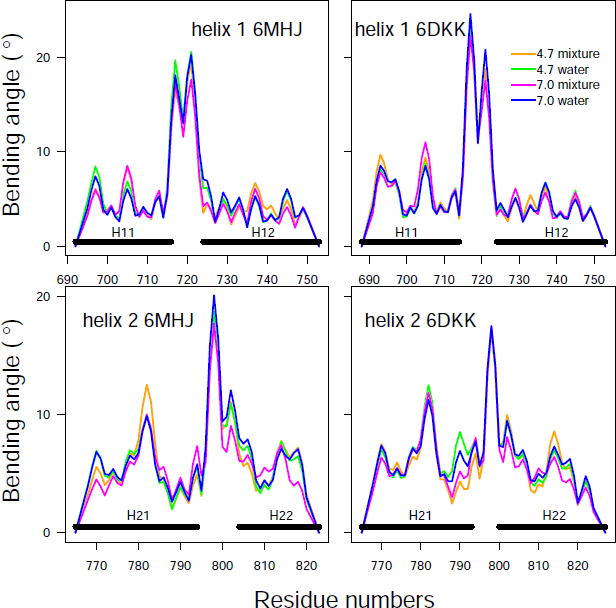
Bending angles of the helices 1 and 2. The analysis was performed using the package Bendix [28] along the first trajectory replica.

We should first stress that the present study intends to predict the behavior of the protein at the approach to the water-membrane interface [10] based on an *in silico* analysis of the protein conformations in solution. In this system, the helices 1 and 2 display long-range interactions with the other parts of the protein structure, which makes the analysis of the hydrophobic interactions very complex. It is thus not simple to draw a conclusion about the *global* equilibrium of the torques, in particular since the membrane is not present explicitly. Nevertheless, we can calculate a total torque acting on each helix part assuming that each part can be modeled as a cylindrical tube. This model is valid in the first approximation since the local bending angle is small within each part (see again Fig. 7). We additionally make the (very strong) assumption that the helix bundle at first makes contact with the membrane in the middle between the helix parts, close to the protruding switch. This assumption is supported by the mutagenesis done by Lam *et al* [8]. This implies that the hydrophobic residues at the boundary of each helix part (bold italic in Table S3) are the ones which enter the membrane first, thereby defining the direction of the resulting hydrophobic strip on the helices. The total torque is thus obtained by summing up the local torques *T̃_kl_* along each region H11, H12, H21 and H22, relative to these residues.

The resulting total torques fluctuate only marginally around a constant value along the trajectory time (Figs 8 and S11, S12). The type of solvent does not induce much variations in the torque values. Changing the protonation scheme slightly shifts the value for H22, since one additional residue (GLU-809) gets protonated at pH=4.7. The largest difference is observed between 6MHJ and 6DKK structures: the value for H22 is negative and close to zero for 6MHJ and positive for 6DKK. Due to the existence of the coiled-coil motifs we assume a direct interaction of the region H11 with H22 and of the region H12 with H21, respectively. For the local torques to act on the membrane, the global torque has to vanish. This is approximately the case for the conformation 6DKK at pH=4.7, for both coiled-coil motifs. Interestingly, this is the structure obtained at conditions (acidic pH and protruding switch) pointed out [8] as characteristic for the interaction with the membrane.

**Figure 8:**
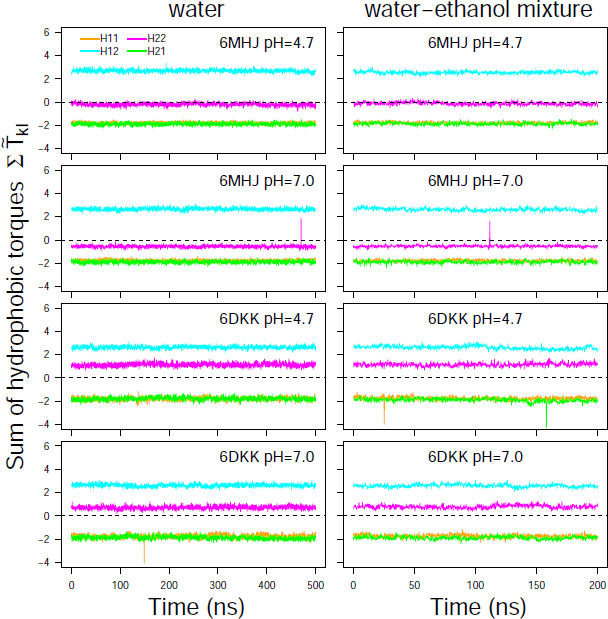
Torque values averaged along the residues 695-713 (H11: orange), 730-748 (H12: cyan), 766-784 (H21: green) and 801-819 (H22: magenta) and plotted along the trajectory time. H12 and H21 are interacting through a coiled-coil motif, as well as H11 and H22. For pH 7, these values were respectively calculated on the following sets of hydrophobic residues: I695, A698, L699, W706, V709, I713 for H11, M732, A735, L736, A740, A742, A745, I746, I747 for H12, I766, L769, L773, I777, A780, M781, I782, I784 for H21, I801, P802, G804, V805, L808, F811, A813, L815, A818, L819 for H22. For pH 4.7, the protonated residue E809 should be added to the list of H22. Note that the curves of H11 and H21 superimpose each other.

The opposite torque values suggest a mechanism in which the helix halves in direct interaction *via* the coiled-coil motifs (H11/H22 and H12/H21, respectively) undergo opposite torques. These global torques could induce additional rotations of each helix with respect to the other, inducing the hydrophobic residues involved in the coiled-coil interactions to insert between the lipid tails of the membrane.

In the structures of the translocation domain studied here, the helices 1 and 2 are surrounded by the N terminal domain, the switch domain, the C terminal *α* helix and the tails connecting the switch and the helix 1 and connecting the helix 2 to the C terminal helix (Table 1). From the previous analyses (Fig S2), the C terminal helix displays a tendency to unfold. In addition, sites of ethanol binding are observed on the tails (Fig S3). Moreover, Lam *et al.* proposed that the *β* hairpin switch anchors the protein to the membrane [8]. Putting all that together, it is quite probable that, at a certain stage of the membrane interaction, the helices 1 and 2 correspond to the last folded part of the translocation domain. At that stage, the torque profile plays thus an essential role for predicting the conformational transition of the two coupled tubes in the membrane.

## 4 Discussion

In this work the conformational dynamics of the BoNT translocation domain was investigated by molecular dynamics simulations. Two different experimental structures have been employed as starting points of the simulations, performed at acidic and neutral pH values. The protein-membrane interaction was implicitly taken into account by using a water-ethanol mixture to mimic a hydrophobic environment. The ethanol displays a polarity of 0.6 on a scale where water is at 1, and hexane at 0 [29]. Ethanol is formed of a polar group (OH) and of a hydrophobic tail (ethane) displaying some similarity with the lipid molecules, formed of a polar head and a tail. In the view of the authors, the water-ethanol mixture provides a numerical trick for testing the hydrophobic interactions of the protein as the ethanol molecules are smaller than the lipids and can interact more freely with the protein surface. The reliability of water-ethanol for mimicking the effect of the membrane is supported by the behavior of the switch in the water-ethanol mixture. In contrast to the observations in aqueous solution, the switch was indeed shown to keep its protruding position in mixed solvent. This also confirms that the protruding conformation is probably the one selected in establishing the first interaction with the membrane.

The present work may also be valuable in the frame of enhanced sampling approaches [30, 31], as a key step in these methods is the choice of collective variables, defining the coordinate of reaction for the exploration of the conformational space of biological systems. In the case of soluble proteins undergoing translocation across the membrane, this choice is very delicate as the distribution of hydrophobic residues on the protein surface cannot give an indication, in contrast to the transmembrane proteins. The procedure described in this work could provide information for the choice of such collective variables. One possible choice would be the angles between the axes of H11 and H12 and between the axes of H21 and H22 to drive rotation around the pivot, and rotation of each of these helix moities around their own axis to favor the exposure of hydrophobic residues. It is worth noting that the use of a model issued from the soft matter field allowed to shine a new light on this problem. We took advantage of the mesoscopic model of filament interactions [19] to mimic the cooperative effect of the numerous lipids present in an ordered way in the membrane. Indeed, the torque produced by the asymmetry of the positions of hydrophobic residues in the protein can only be active in the presence of a formed membrane, stabilized by the cooperativity of the interactions between lipids.

Based on the conformations explored at the atomistic level, the concomitant interaction between the coiled-coil helices and the membrane was investigated in the frame of the mesoscopic *Twister* model, which permits to calculate the torques underlying the membrane-protein interaction, required for membrane opening. The introduction of this model is the most innovative point of the work presented here. Its novelty lies in the connection established between conformations at atomic resolution and the mesoscopic scale. A possible extension would be to use an adapted scale of hydrophobicity for the amino-acid residues. Various scales could be exploited in that respect [32, 33]. Nevertheless, the present state of the model only allows for a qualitative description of protein-membrane interaction. A description at atomic resolution of this interaction would require the development of a related energy function. Moreover, while a lipid-protein complex relies on relatively localized interactions, in principle the ethanol-protein interaction would take place *via* diffuse contacts. However, the analysis of the ethanol distribution around the structure 6DKK at pH 4.7 revealed some strong preferred localisation of ethanol methyl groups close to the center of the translocation domain (large green spots in Fig.5, lower right panel). Knowing that the switch, located at the center of the domain as well, has also some preference for insertion into the membrane, these two pieces of information permit to infer a local pivot point for applying the torques arising from the asymmetry in the positions of the hydrophobic residues.

The implication of coiled-coil motifs in this study is not surprising, since these motifs are known to play a role in many other structures interacting with membranes such as in the SNARE complex [34]. In addition, successful examples in the design of membrane systems used coiled-coil motifs as a starting point. For example, rational *de novo* design of channels was possible based on *α*HB coiled-coil peptides [35]. Moreover, a liposomal fusion model system was developed [36] using a set of hetero-dimeric coiled-coil peptide pairs.

The *Twister* analysis of the torques along the helix tubes permits to propose a model in which local torques induce membrane deformations. The resulting global opposite torques within the coiled-coil motifs of helices 1 and 2 additionally imply a rotation of the motif H11 with respect to H22 and of H12 with H21, respectively. The middle regions of helices 1 and 2 were not included in the torque analysis, since they were not part of a coiled-coil motif. As these regions are close to the *β* hairpin switch, their unfolding could, however, permit a reorganisation of the four helix motifs, potentially inducing the formation of a pore in the membrane. Interestingly, the beltless translocation domain of botulinum toxin A was shown to be able to act as a conductive channel [37], reminiscent of the design experiments on coiled-coil motifs [35, 36].

The analysis of the trajectories at various pH values and in polar and more hydrophobic solvents allows to propose hypotheses on the interactions between the *whole* botulinum toxin (BoNT) and the membrane of neurotransmitter vesicles. Firstly, upon acidification the translocation domain adopts a conformation with protruding hairpin to initiate the interaction with the membrane. Secondly, the unfolding of the C-terminal *α* helix agrees with the observations made on the BoNT/E1 structure [6]: the C-terminal *α* helix (residues 821-845 in BoNT/E1) of the translocation domain unfolds and displays loop conformations. The translocation domain behaves thus relatively independently from the remaining BoNT structure, in agreement with the modularity previously observed between various regions of BoNT [7]. The most stable sub-structures of the translocation domain are the coiled-coil motifs formed between helices 1 and 2; these motifs apply torques opening the membrane, possibly inducing a pore formation. Several models of the subsequent passage of the catalytic domain of BoNT have been proposed [38]. An investigation of the actual translocation mechanism is out of the scope of the present study. In summary, this study represents a first step of coupling the mesoscopic *Twister* model to an atomistic approach with predictive power for protein-membrane interactions. The concomitant use of the two approaches allowed to define a hierarchy in the interactions between the translocation domain and the membrane. This hierarchy proposes a division of the protein structure in regions undergoing unfolding or separation from the protein core, and the bundle of helices 1 and 2 as the main player involved in membrane insertion. Our predictions are of particular interest in the case of soluble proteins displaying conformational changes during their insertion in the membrane. As structural biology techniques often have difficulties to capture these changes due to the dynamics of the process, the model described here might provide low resolution information which can be probed experimentally or combined with high resolution structures.

## 5 Materials and Methods

### 5.1 Preparation of the starting conformations of toxins

The monomeric (6MHJ) and dimeric (6DKK) structures of the translocation domain [8] have been downloaded from the Protein Data Bank www.rcsb.org. In 6MHJ, the residue F658 has been modified into E658, in order to fit to the WT sequence of BoNT/A1. In 6DKK, the chain B has been extracted as a monomer, as it was containing the smallest number of missing residues at the C terminal extremity of the protein. As already mentioned in the introduction, a 6DKK monomer was studied by molecular dynamics as the dimer was shown [8] to be induced by crystallogenesis.

The studied systems were named (Table S1) using the PDB entry name (6MHJ/6DKK), the pH for which the protonation was defined (47 or 70) and the solvent name (W for TIP3P and E for water-ethanol). The names of these systems will also be the names of the corresponding trajectories.

### 5.2 Molecular dynamics simulations

For each previously described system, the protein was embedded in a water box (Table S1), and counterions were added to neutralize the net system charge. The total number of atoms varies between 97483 and 189355 (Table S1). All molecular dynamics (MD) simulations were performed using NAMD 2.13 [39] and NAMD 3.0 [40] depending on the type of machine architecture. The CHARMM36 force field [41] for the protein and the TIP3P model for the water molecules [42] were used. A cutoff of 12 Å and a switching distance of 10 Å was employed for non-bonded interactions, while long-range electrostatic interactions were calculated with the Particle Mesh Ewald (PME) method [43]. Before starting each MD trajectory, the system was minimized for 20000 steps, then heated up gradually from 0 to 310 K in 31000 integration steps. Finally, the system was equilibrated for 500.000 steps in the NVT ensemble at 310 K, yielding a trajectory of 1 ns. In these first three stages, all carbon *α* (C*α*) atoms were kept fixed. Simulations were then performed in the NPT ensemble, (*P* = 1 bar, *T* = 310 K), with all atoms free to move. During the equilibration and production trajectories, the non-iterative Settle algorithm [44, 45] was used to keep rigid all covalent bonds involving hydrogen atoms, enabling a time step of 2 fs. Atomic coordinates were saved every 20 ps. 500 ns of production were recorded and each trajectory was triplicated for a cumulative trajectory duration of 1.5 *µ*s.

The 50/50 mixed water-ethanol systems were prepared in the following way. The software PACKMOL [46] was run to generate boxes of water and ethanol molecules, with the same number of each molecule type (water-ethanol box in Table S1). This box was then minimized, thermalized and equilibrated for 1 ns using the same protocol as the one described above for the protein in water. The final frame was then superimposed to the protein’s initial conformations solvated in a water box, and each water or ethanol molecule located at less than 2.4 Å from the protein atoms was discarded from the system. The NAMD coordinate and topology input files were generated and the minimization, thermalization, equilibration and production protocol described above was used to simulate the protein-ethanol-water systems. 200 ns of production were recorded and each trajectory was triplicated for a cumulative trajectory duration of 0.6 *µ*s. For the trajectories recorded on 6DKK, roto-translational motions of the protein were harmonically restrained throughout the simulation (with a scaled force constant of 30 kcal/mol) using the colvar commands[47], to avoid any rigid-body motion of the protein but leaving unaltered the internal dynamics.

### 5.3 Analysis of the atomistic trajectories

The analysis of protein conformations sampled along MD trajectories was realized using ccptraj [48] and the python package MDAnalysis [49, 50]. The bending angles of helices 1 and 2 (Fig. 1A) in H*_N_* were determined using the VMD plugin Bendix [28] on every 10 frames of the trajectory.

Due to the elongated structure of the translocation domain, the whole shape of the domain was analyzed using a model based on a cylinder. Using the MDAnalysis library [50], the moments of inertia and principal axes were calculated from the positions of the C*α* atoms, on every 10 frames of the trajectory. With the help of the moment of inertia and the principal axes of the translocation domain, the average radius *r* of a cylinder superposed to the protein was calculated as: 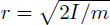, where *I* is the moment of inertia with respect to the axis of the cylinder and *m* is the total molar mass of the C*α* atoms.

To analyze the solvent partition in the protein domain, the spatial density function (SDF) distribution of the solvent molecules around the protein was calculated, using the VolMap tool provided by VMD [51]. After translational and rotational fitting of the system to a reference conformation of the protein (the one in the initial frame), the positions of selected solvent atoms (the oxygen atom in water, the oxygen atom of the hydroxyl moiety and the carbon atom of the methyl group in the ethanol, respectively) were mapped onto a cubic grid with a grid spacing of 1 Å centered on the protein. The grid densities have then been averaged over the number of analyzed frames.

### 5.4 Coarse-grained modeling of the helices 1 and 2

The helices 1 and 2 (Fig. 1A) were described in a coarse-grained manner: we have assumed that they are elastic cylindrical tubes. In that framework, the axes of helix 1 and helix 2 were defined using the method originally developed by Kahn [52]. The starting and last protein residues of the tubes were fixed using the helix limits given in Table 1. The hydrophobic residues considered in a given *α* helix were selected in the following way: the residues GLY, ALA, VAL, LEU, ILE, PRO, PHE, MET and TRP classically considered as hydrophobic, supplemented at pH 4.7 with the protonated residue GLU-809. Each tube could interact with the membrane *via* the hydrophobic attraction between the protein residues and the lipid tails of the membrane bilayer. To understand this interaction, we had to consider the orientations of the residues with respect to each other.

Using the relative positions of hydrophobic residues *k* of the considered *α* helix (Table S3), a set of vectors was determined which are normal to the axis. Each vector *V_k_* connects the axis to the C*α* atom of residue *k* (see Fig 9). If all of these vectors were parallel, the hydrophobic residues would align on a straight line, the helix would be amphipathic and could insert itself into the membrane without undergoing an elastic deformation. When the angle *θ_kl_* between *V_k_* and *V_l_* of two consecutive hydrophobic residues *k* and *l* is non zero, the tube has to twist to align the two residues. The torque *T_kl_* resulting from the non-alignment of the residue is proportional to the angle *θ_kl_* in lowest order and anti-proportional to the distance *D_kl_* between the points *P_k_* and *P_l_*. For a cylindrical tube *T_kl_* is given by [53, 54, 55]

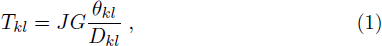

**Figure 9:**
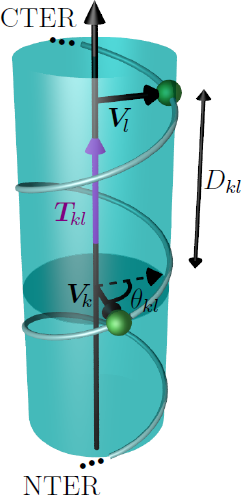
Geometry of the coarse-grained *α* helix describing the calculation of the torque *T_kl_* (in purple). The *α* helix is represented by a transparent cylinder, the helix axis is parallel to the torque. The two vectors ***V*** *_k_* and ***V*** *_l_* are perpendicular to the helix axis and connect the axis to the C*α* atoms of the two consecutive hydrophobic residues *k* and *l* (in green). The projection of *V_l_* next to *V_k_* (dashed vector) is drawn in order to illustrate the definition of *θ_kl_*.

where 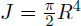 is the polar moment of inertia of the tube’s cylindrical cross section with radius *R* and *G* is the shear modulus. In the following we will use the scaled torque 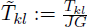. For proteins, due to the variability of their structures, only sparse experimental information is available for the value of *GJ*. Indeed, only few cases have been studied in detail: values for a large protein complex (IgG/G protein) are of the order of 45000 pNnm^2^, whereas a value of only 400 pNnm^2^ was found for the flexible DNA double helix [54, 55]. From the point of view of sizes and internal mobility the BoNT helices 1 and 2 are in between these two cases. We can thus expect values between the numerical values quoted above.

The torque vectors are all parallel to the axis. A positive sign of *T̃_kl_* indicates that the vector is aligned from the N to the C terminal extremities of the helix sequence, whereas it is aligned in the opposite direction when the sign is negative. A python script using the library MDAnalysis [50] has been specifically developed for the calculation of the torques.

## 5 Author contributions

Conceptualization, T.E.M. and M.M.M.; methodology, G.C., T.E.M. and M.M.M.; investigation, A.D., G.C. and T.E.M.; writing—original draft preparation, G.C.; writing—review and editing, T.E.M. and M.M.M.; visualization, A.D.; supervision, M.M.M.; funding acquisition, T.E.M. All authors have read and agreed to the published version of the manuscript.

## 6 Funding

The authors thank the GENCI (project A0080710764) for supercomputing time. TEM and MMM acknowledge CNRS and University of Lorraine for financial support. GC acknowledges the University of Palermo (grant FFR-Cottone).

## 7 Data availability

The dataset(s) supporting the conclusions of this article is(are) available at zenodo.org. The exact address at zenodo will be determined at the step of manuscript revision.

The authors declare no conflict of interest.

## Notes

### Competing Interest Statement

The authors have declared no competing interest.

