## Supplementary Materials for "Conformational Space of the Translocation Domain of Botulinum Toxin: Atomistic Modeling and Mesoscopic Description of the Coiled-Coil Helix Bundle"

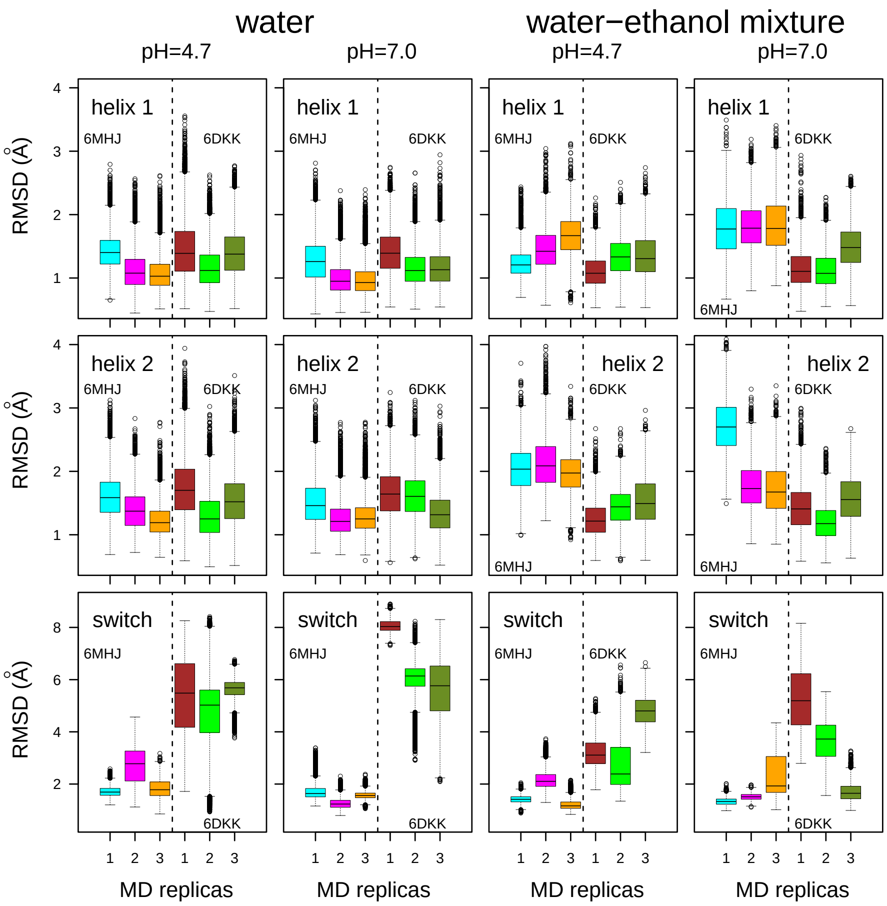


Figure S1: Distributions of coordinate RMSD (Å) for specific domains of the protein.

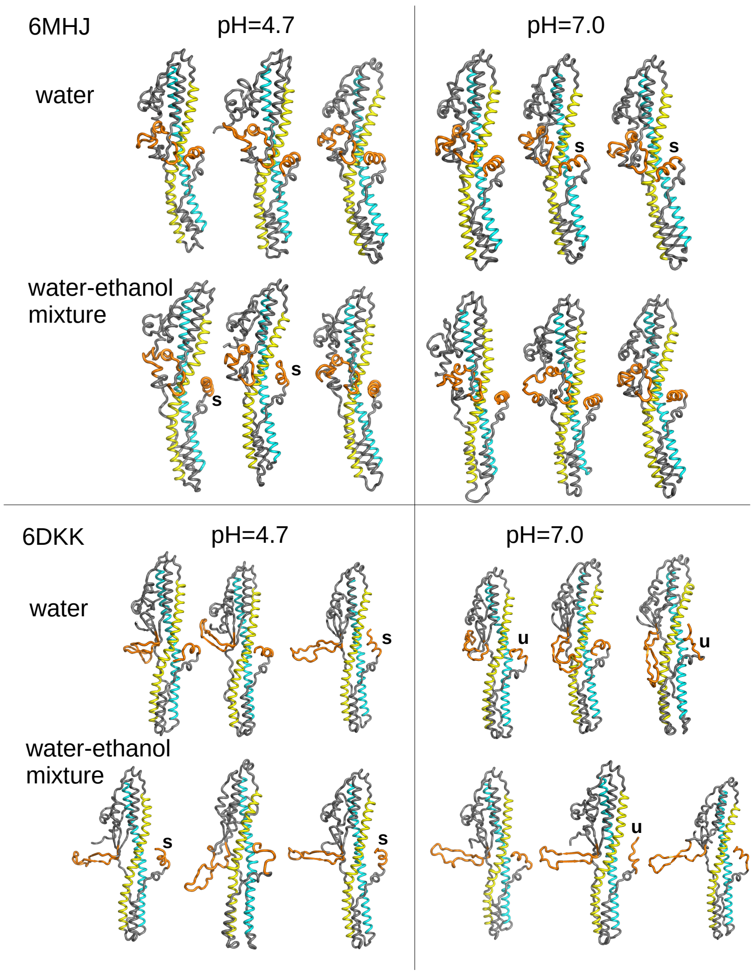


Figure S2: Final conformations of the three replica trajectories in each investigated system. The protein is represented in ribbon; the switch region and the C terminal $\alpha$ helix are colored in orange and the helices 1 and 2 in cyan and yellow, respectively. Examples of unfolded C terminal $\alpha$ helices are labeled with 'u', and examples of different orientations of the C terminal $\alpha$ helices are labeled with 's'.

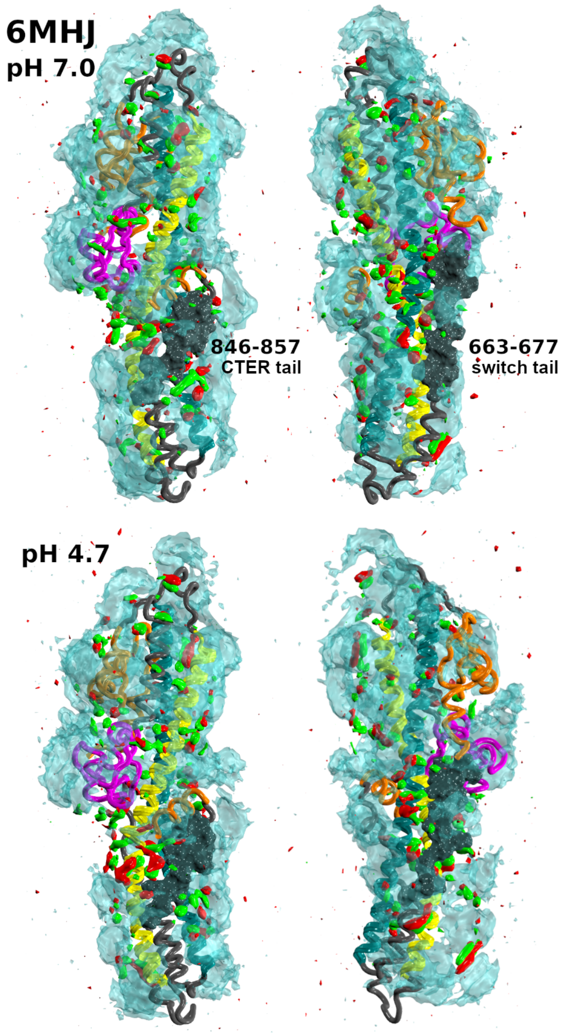


Figure S3: Isosurfaces of the Spatial Density Function of water and ethanol around the protein, represented as in Figure 6 in the main text. The isodensity value is 0.0115 for all atomic species. The protein regions are colored as in Figure 1. The residue stretches 663-677 and 846-857 (switch and CTER tails, respectively) are highlighted in dark grey surface representation. Data were collected from the three replica trajectories. This image was prepared using VMD [1].

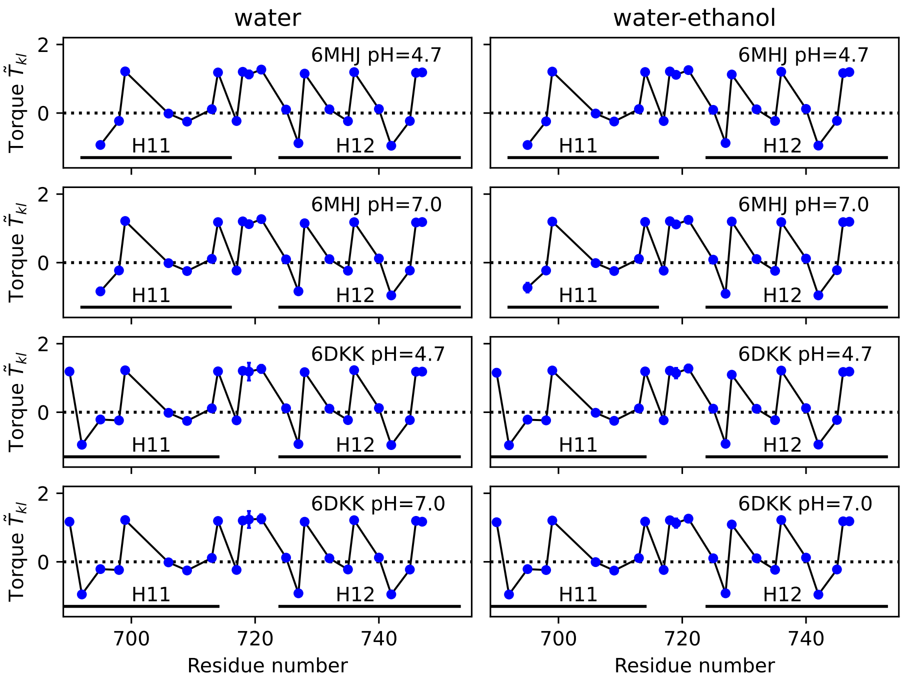


Figure S4: Local torque values $\tilde{T}_{kl}$ along the helix 1. Values averaged along the second trajectory replica and their standard deviations are plotted as a function of the residue number. Standard deviations are smaller than the size of the data points except for residue ALA-719.

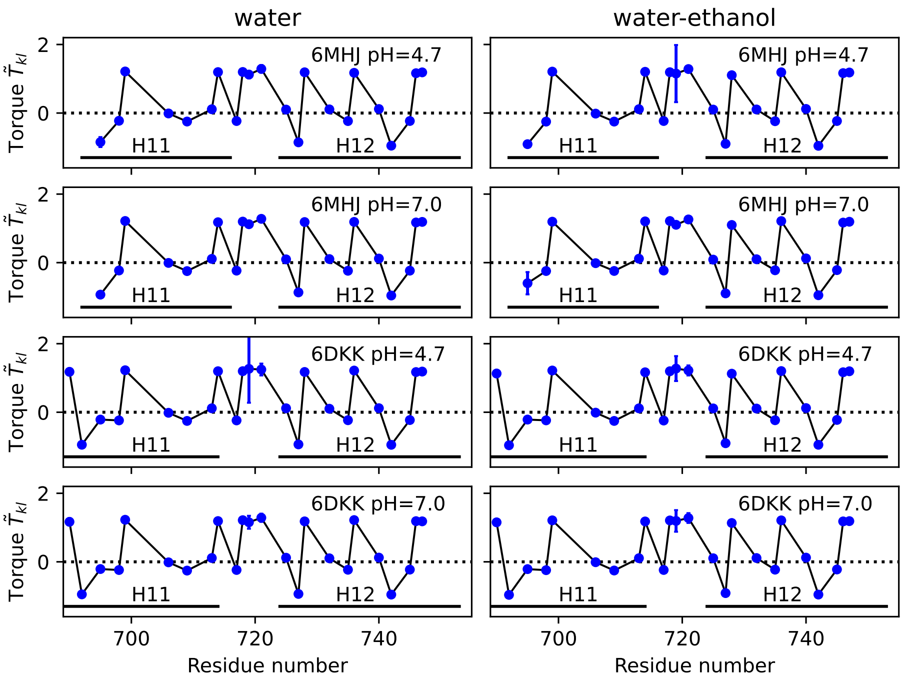


Figure S5: Local torque values $\tilde{T}_{kl}$ along the helix 1. Values averaged along the third trajectory replica and their standard deviations are plotted as a function of the residue number. Standard deviations are smaller than the size of the data points except for residue ALA-719.

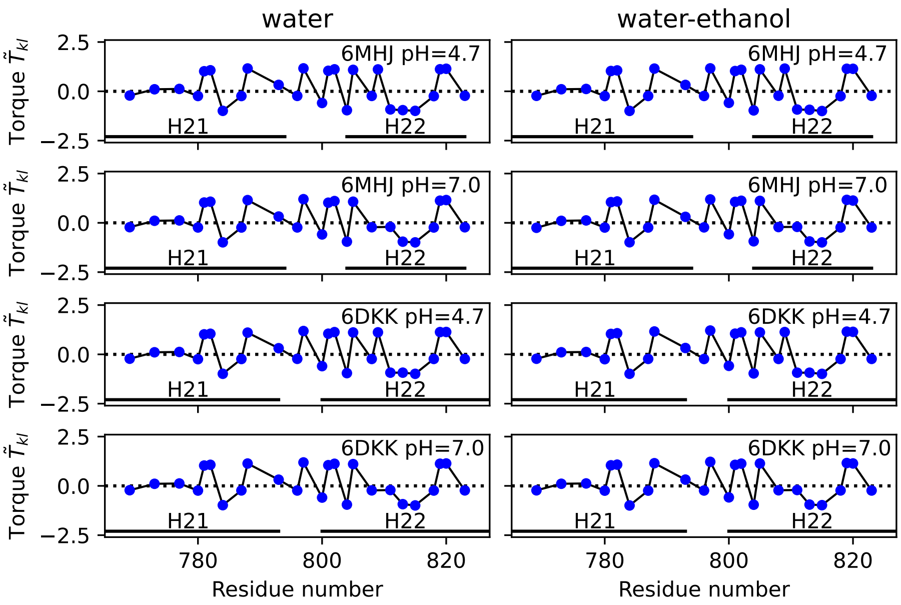


Figure S6: Local torque values $\tilde{T}_{kl}$ along the helix 2. Values averaged along the first trajectory replica and their standard deviations are plotted as a function of the residue number. Standard deviations are smaller than the size of the data points.

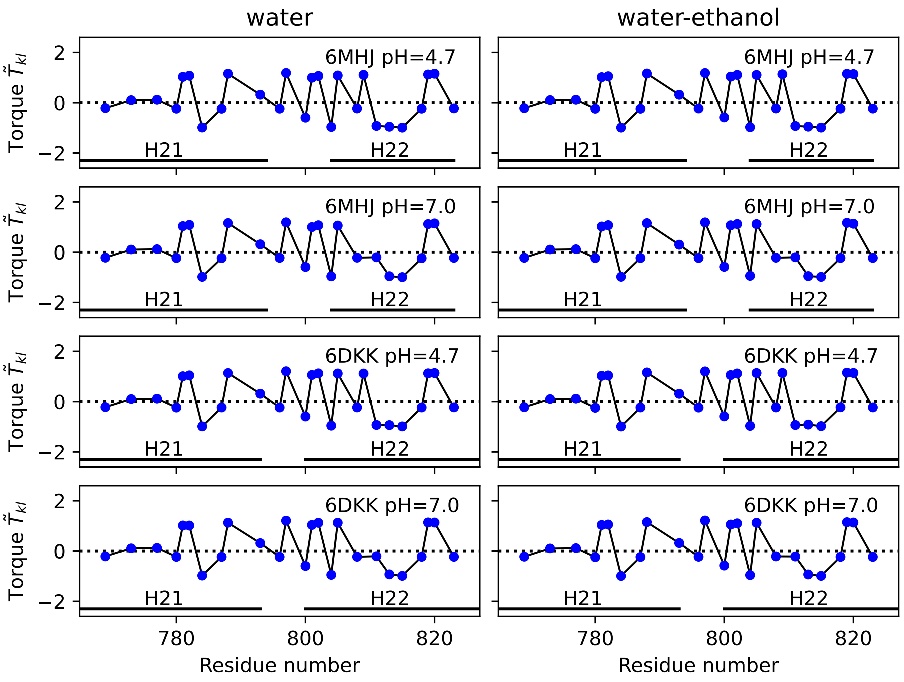


Figure S7: Local torque values $\tilde{T}_{kl}$ along the helix 2. Values averaged along the second trajectory replica and their standard deviations are plotted as a function of the residue number. Standard deviations are smaller than the size of the data points.

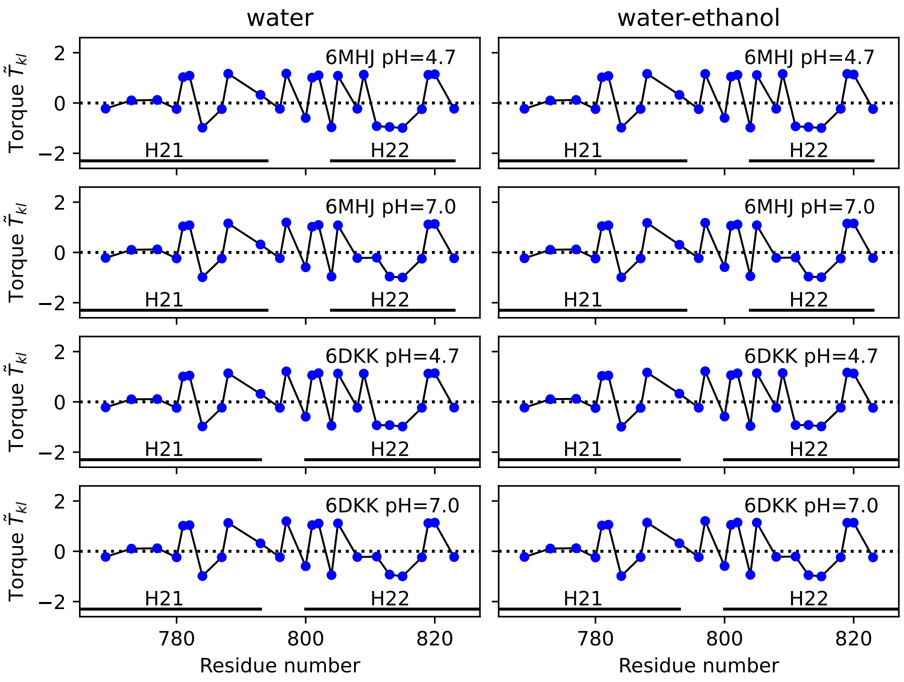


Figure S8: Local torque values $\tilde{T}_{kl}$ along the helix 2. Values averaged along the third trajectory replica and their standard deviations are plotted as a function of the residue number. Standard deviations are smaller than the size of the data points.

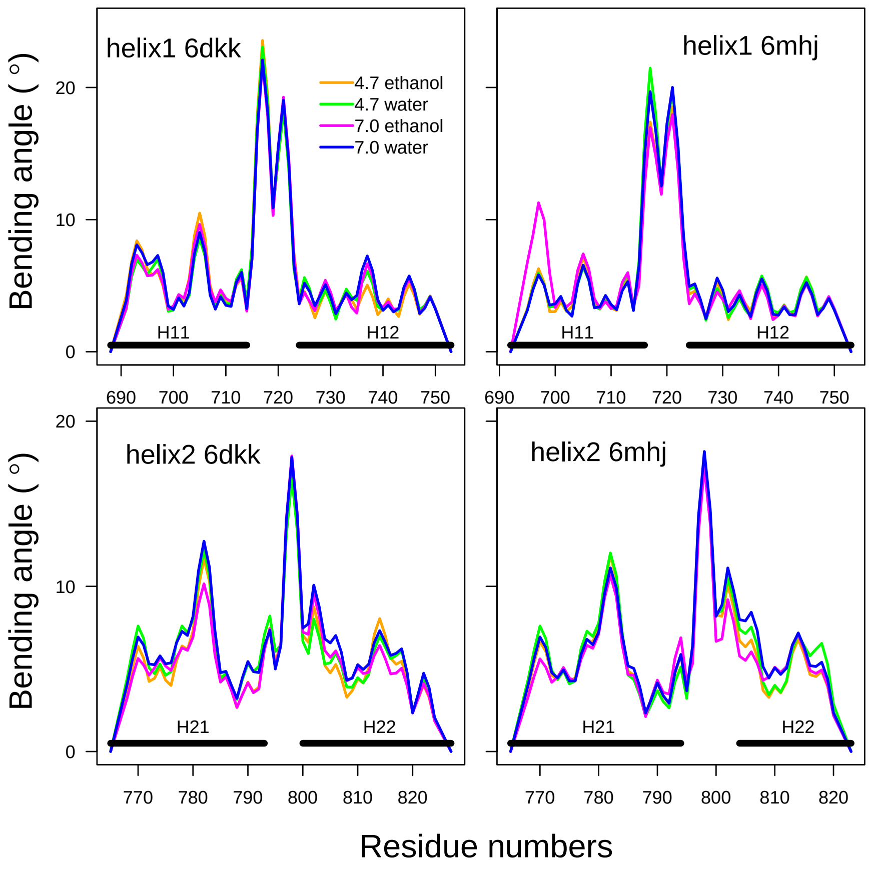


Figure S9: Bending angles of the helices 1 and 2. The analysis was performed using the package Bendix [2] on the second trajectory replica.

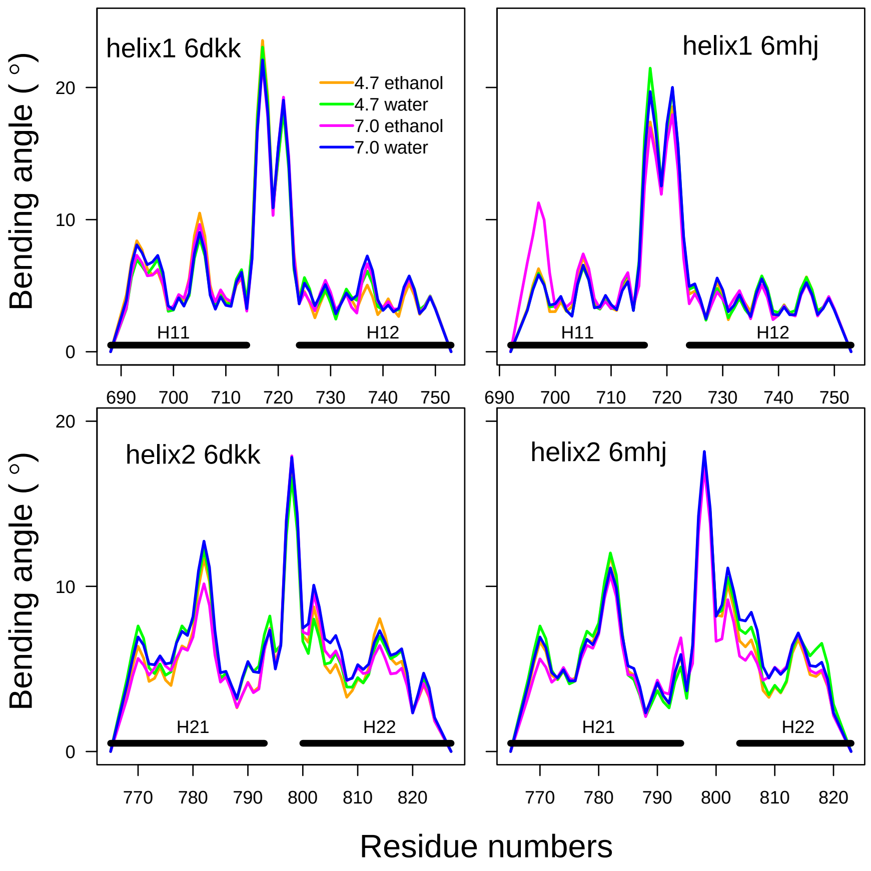


Figure S10: Bending angles of the helices 1 and 2. The analysis was performed using the package Bendix [2] on the third trajectory replica.

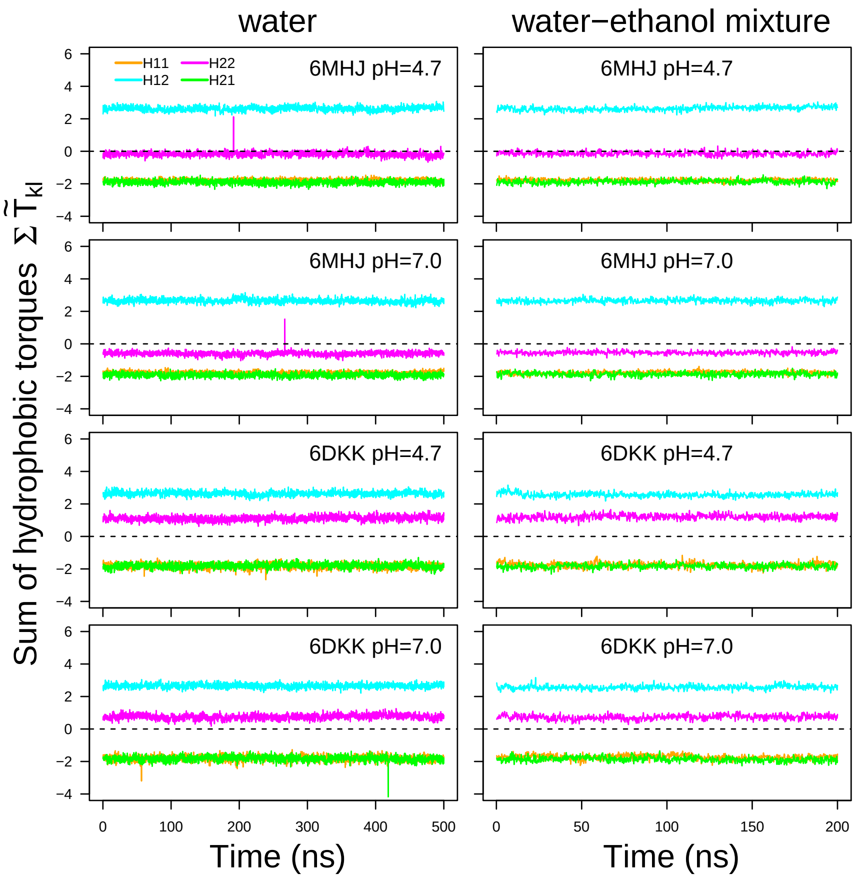


Figure S11: Torque values averaged along the residues 695-713 (H11: orange), 730-748 (H12: cyan), 766-784 (H21: green) and 801-819 (H22: magenta) and plotted along the time for the second trajectory replica. H12 and H21 are interacting through a coiled-coil motif, as well as H11 and H22. For pH 7, these values were, respectively, calculated on the following sets of hydrophobic residues: I695, A698, L699, W706, V709, I713 for H11, M732, A735, L736, A740, A742, A745, I746, I747 for H12, I766, L769, L773, I777, A780, M781, I782, I784 for H21, I801, P802, G804, V805, L808, F811, A813, L815, A818, L819 for H22. For pH 4.7, the protonated residue E809 was added to the list of H22. Note that the curves of H11 and H21 superimpose each other.

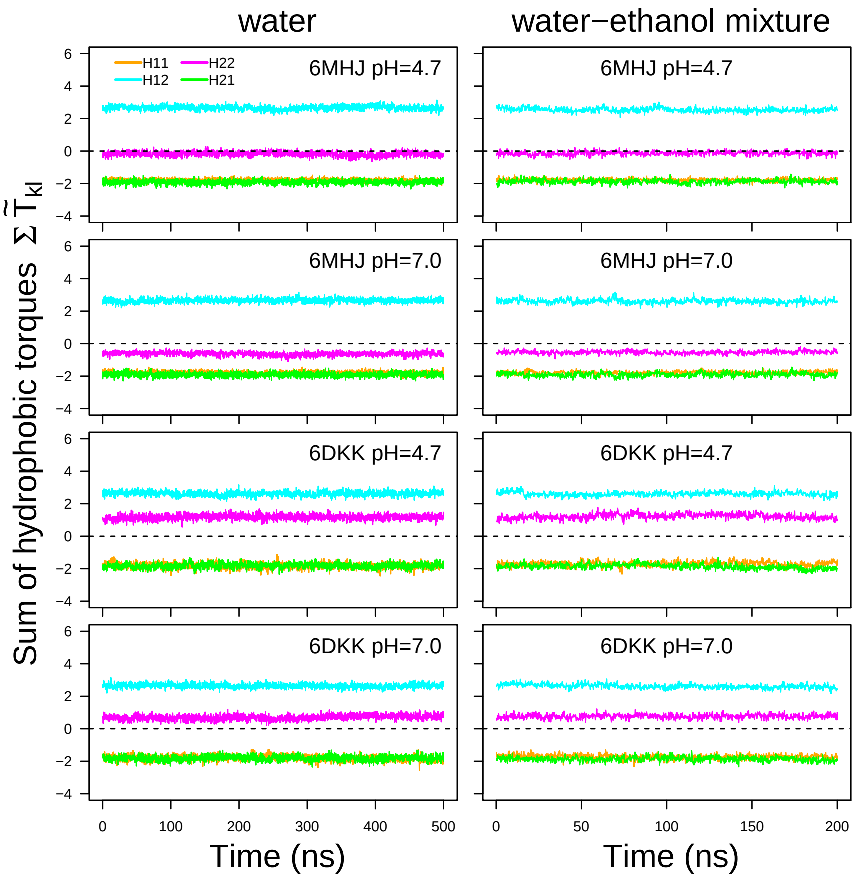


Figure S12: Torque values averaged along the residues 695-713 (H11: orange), 730-748 (H12: cyan), 766-784 (H21: green) and 801-819 (H22: magenta) and plotted along the time for the third trajectory replica. H12 and H21 are interacting through a coiled-coil motif, as well as H11 and H22. For pH 7, these values were, respectively, calculated on the following sets of hydrophobic residues: I695, A698, L699, W706, V709, I713 for H11, M732, A735, L736, A740, A742, A745, I746, I747 for H12, I766, L769, L773, I777, A780, M781, I782, I784 for H21, I801, P802, G804, V805, L808, F811, A813, L815, A818, L819 for H22. For pH 4.7, the protonated residue E809 was added to the list of H22. Note that the curves of H11 and H21 superimpose each other.

| System | number of water molecules | number of ethanol molecules | counter-ions | total number of atoms | solvent box (Å,Å,Å) |
|---|---|---|---|---|---|
| 6MHJ47W | 30728 | - | 6 Cl- | 97483 | 146.8, 82.7, 79.8 |
| 6MHJ47E | 15544 | 15270 | 5 Cl- | 189355 | 187.6, 103.6, 102.2 |
| 6MHJ70W | 30734 | - | 2 Na+ | 97489 | 146.7, 82.7, 79.8 |
| 6MHJ70E | 15542 | 15274 | 3 Na+ | 189375 | 187.6, 103.6, 102.3 |
| 6DKK47W | 39954 | - | 3 Cl- | 125155 | 127.2, 113.6, 86.3 |
| 6DKK47E | 15195 | 15029 | 3 Cl- | 186139 | 183.9, 104.1, 102.2 |
| 6DKK70W | 39957 | - | 2 Na+ | 125157 | 127.1, 113.5, 86.2 |
| 6DKK70E | 15201 | 15033 | 2 Na+ | 186186 | 184.0, 104.1, 102.2 |
| water-ethanol box | 16120 | 16120 | - | 193440 | 192.1, 104.1, 101.8 |

Table S1: Composition of the systems for molecular dynamics simulations. The box sizes are those observed at the end of equilibration.

| PDB entry | Residues | RMSD (Å) | $\omega_0$ (deg) | $\alpha$ (deg) | Rise (Å) |
|---|---|---|---|---|---|
| 6MHJ | 695-713/801-819 | 0.643 | 1.636 | 5.958 | 1.503 |
| 6MHJ | 730-748/766-784 | 0.472 | 3.644 | 13.019 | 1.514 |
| 6DKK | 695-713/801-819 | 0.59 | 1.742 | 6.465 | 1.497 |
| 6DKK | 730-748/766-784 | 0.579 | 3.881 | 14.013 | 1.498 |

Table S2: Crick parameters determined on the residues ranges extracted from PDB entries 6MHJ and 6DKK, using the software CCCP (Coiled-coil Crick Parameterization) available at the Web server: `www.grigoryanlab.org/cccp` [3]. For 6DKK, the calculation was performed on the chain A. The RMSD value is calculated between the PDB structure and the ideal geometry determined using the measured Crick parameters. Some Crick parameters are given in the Table: Superhelical frequency ($\omega_0$), pitch angle ($\alpha$), rise per residue (Rise).

| PDB entry | pH | helix | Number of residues | residue list |
|---|---|---|---|---|
| 6MHJ | 4.7 | helix 1 | 23 | 692, 695, 698, 699, 706, 709, 713, ***714***, 717, 718, 719, 721, ***725***, 727, 728, 732, 735, 736, 740, 742, 745, 746, 747 |
| 6MHJ | 4.7 | helix 2 | 27 | 766, 769, 773, 777, 780, 781, 782, 784, 787, 788, ***793***, 796, 797, 800, 801, 802, ***804***, 805, 808, **809**, 811, 813, 815, 818, 819, 820, 823 |
| 6MHJ | 7.0 | helix 1 | 23 | 692, 695, 698, 699, 706, 709, 713, ***714***, 717, 718, 719, 721, ***725***, 727, 728, 732, 735, 736, 740, 742, 745, 746, 747 |
| 6MHJ | 7.0 | helix 2 | 26 | 766, 769, 773, 777, 780, 781, 782, 784, 787, 788, ***793***, 796, 797, 800, 801, 802, ***804***, 805, 808, 811, 813, 815, 818, 819, 820, 823 |
| 6DKK | 4.7 | helix 1 | 25 | 689, 690, 692, 695, 698, 699, 706, 709, 713, ***714***, 717, 718, 719, 721, ***725***, 727, 728, 732, 735, 736, 740, 742, 745, 746, 747 |
| 6DKK | 4.7 | helix 2 | 27 | 766, 769, 773, 777, 780, 781, 782, 784, 787, 788, ***793***, 796, 797, ***800***, 801, 802, 804, 805, 808, **809**, 811, 813, 815, 818, 819, 820, 823 |
| 6DKK | 7.0 | helix 1 | 25 | 689, 690, 692, 695, 698, 699, 706, 709, 713, ***714***, 717, 718, 719, 721, ***725***, 727, 728, 732, 735, 736, 740, 742, 745, 746, 747 |
| 6DKK | 7.0 | helix 2 | 26 | 766, 769, 773, 777, 780, 781, 782, 784, 787, 788, ***793***, 796, 797, ***800***, 801, 802, 804, 805, 808, 811, 813, 815, 818, 819, 820, 823 |

Table S3: Hydrophobic residues for helices 1 and 2 depending on the PDB structures and pH values. The residue 809, which is protonated at pH 4.7, is written in bold. The residues defining the direction of the resulting hydrophobic strip on the helices are written in bold italic, as described in Sec. III.C in the main text.
